# The power of mixed survey methodologies for detecting decline of the Bornean orangutan

**DOI:** 10.1101/775064

**Authors:** Truly Santika, Kerrie A. Wilson, Erik Meijaard, Marc Ancrenaz

## Abstract

For many threatened species, it is difficult to assess precisely for large areas the change in their abundances over time and the relative impacts of climate and anthropogenic land use. This is because surveys of such species are typically restricted to small geographic areas, are conducted during short time periods, and use different survey protocols. We assessed the change in the abundance of Bornean orangutan *Pongo pygmaeus morio* in Sabah, Malaysia, and to identify environmental drivers affecting the change by integrating different types of survey data. We used nest count data obtained from aerial and ground transect surveys and occurrence data obtained from reconnaissance walks and interview survey over the past decade. We built a spatially-explicit dynamic population model within the Bayesian framework allowing these varying survey data to be analyzed jointly by explicitly accounting for each survey’s sampling rate. We found that sampling rates vary across survey types, reflecting each survey’s associated effort. Orangutan survival rates were strongly determined by natural forest extent and moderately by temperature. Orangutan migration rates across more than 1 km distance between forest patches were low, which underlines the importance of maintaining ecological connectivity. The paucity of species abundance data collected in a consistent manner over many years across broad extents often hinders the assessment of species population trend and their persistence across regional scales. We demonstrate that this can be addressed by integrating multiple survey data across different localities, provided that sampling rate inherent to each survey is accounted for.

## 1 INTRODUCTION

The development of comprehensive strategies for threatened species management and recovery necessitates an understanding of the impacts of environmental change on species persistence (Estrada & Arroyo 2012). The impacts of environmental change would ideally be assessed using data collected in a consistent manner across the entire extent of interest and over many years. Such data are often not readily available or are costly to obtain, and this is particularly the case for threatened large-ranging species in remote areas (de Fraga *et al*. 2014), such as the Bornean orangutan *Pongo pygmaeus*.

Contemporary anthropogenic factors have accelerated the decline of the Bornean orangutan over the last centuries. Goossens *et al.* (2006) indicated a drastic decline of orangutan numbers in the Lower Kinabatangan region of the Malaysian state of Sabah over the past centuries. Meijaard *et al*. (2010) showed that orangutan encounter rates have been declining since the mid-19th century even in parts of Borneo that remain forested, suggesting that the species once occurred on the island at higher densities. Hunting for meat and wildlife trade and habitat loss due to conversion of forest to agriculture have been the primary threats to the species (Meijaard *et al*. 2011a; Abram *et al.* 2015). The rapid expansion of oil palm plantations has recently posed additional pressure, increasing human-wildlife conflict and the rates of killing of individuals (Davis *et al*. 2013).

Orangutan population surveys present numerous logistical challenges, because individuals exhibit cryptic behavior and generally occur at relatively low densities, leading to very low encounter rates in the forest. Instead of using direct counts of individuals, counts of orangutan sleeping platforms or “nests” are typically used to estimate population densities (Ancrenaz *et al*. 2004b; Kühl 2007), and a diverse range of survey protocols are employed for this purpose. Line transect surveys of orangutan nests are the most commonly employed method (Ancrenaz *et al.* 2004b; Johnson *et al*. 2005). Aerial surveys of orangutan nests via helicopter are less common due to the associated cost and aircraft availability (Ancrenaz *et al.* 2005, 2010), although advances in drone technology allows aerial surveys to be conducted more economically (Wich *et al.* 2015). Interview surveys to ascertain orangutan sightings have also provided an economical approach for assessing the status of orangutan populations, although they are subject to an array of biases associated with expert data (Meijaard *et al.* 2011b). Different survey methods have varying detection rates of orangutans and their nests. For example, survey methodologies that are costly may target areas for surveys where orangutan populations are known to exist, enhancing the detection rate. In contrast, a comparatively cheaper survey method may be conducted more extensively even in places without prior reports of orangutans, thus detection rates based on this type of survey are likely to be lower. Ignoring the detection rates associated with different survey methods can potentially mask the actual distribution and abundance of a species (Gu & Swihart 2004; Kéry & Royle 2010).

There have been numerous studies of the biogeography of orangutan derived by extrapolating known occurrences to un-surveyed locations. This is achieved either by generating density estimates based on orangutan nest counts via Distance sampling methods (Buckland *et al*. 2005) (e.g. Knop *et al*. 2004; Ancrenaz *et al*. 2005, 2010; Johnson *et al*. 2005; Wich *et al*. 2008) or by linking observation data or nest density estimates with a suite of environmental predictors via static distribution modelling techniques (e.g. Gregory *et al*. 2012; Wich *et al*. 2012; Struebig *et al.* 2015). However, spatial and temporal projections using nest density estimates have so far overlooked the dynamic nature of orangutan nest construction. Nest decay rates has been shown to vary spatially depending on forest type and altitude and the rate of nest production is determined by the level of forest disturbance, e.g. between logged and unlogged forest (Mathewson *et al*. 2008).

Given these challenges, assessing the change in the abundance of orangutans over large spatial extents requires a modelling approach that can (a) project the density of orangutan based on nest counts, (b) employ multiple types of survey data, and (c) explicitly account for the detection rates of nests inherent in each survey methodology. A dynamic population model within a hierarchical Bayesian framework is appropriate for this purpose. Indeed, in recent years, the hierarchical Bayesian approach has revolutionized methodology for analyzing ecological data, allowing different types of data to be analyzed jointly under a single framework (Royle & Dorazio 2008; Abadi *et al*. 2010). The approach partitions the model into two broad states: the latent (true) population status and observation of the true status. Each state is governed by different processes (explained through sets of covariates), and they are linked through a predefined mechanism based on expert consensus. The approach has gained much interest, especially in species distribution modeling, because it allows measurement or observation error processes underlying most of the species occurrence and abundance data to be taken into account (Kéry & Royle 2008). Applications of this method are continuously expanding, from providing a static snapshot of species distributions (Kéry & Royle 2008), to more complex models that account for the change in landscapes through time (Yamaura *et al*. 2011; Santika *et al.* 2014).

We assessed the abundance and distribution of the Bornean orangutan over time and space, and determined the contribution of climate and land use dynamics to the observed changes. We focused on the subspecies *P. p. morio* in Sabah, Malaysia, with the aim to inform the management of this species at the level of the entire state (Figure 1a). Unlike the neighboring country of Indonesia where government policy for threatened species management is developed at a national level, state government plays the most important role for regional decision making in Malaysia (Sabah Wildlife Department 2011). We used nest count data obtained from aerial and line transect surveys, presence-absence data obtained from reconnaissance walks, and direct orangutan sightings from interview surveys, over the past decade and developed a dynamic abundance model for the orangutan by integrating data obtained from multiple survey methods.

**Figure 1.**
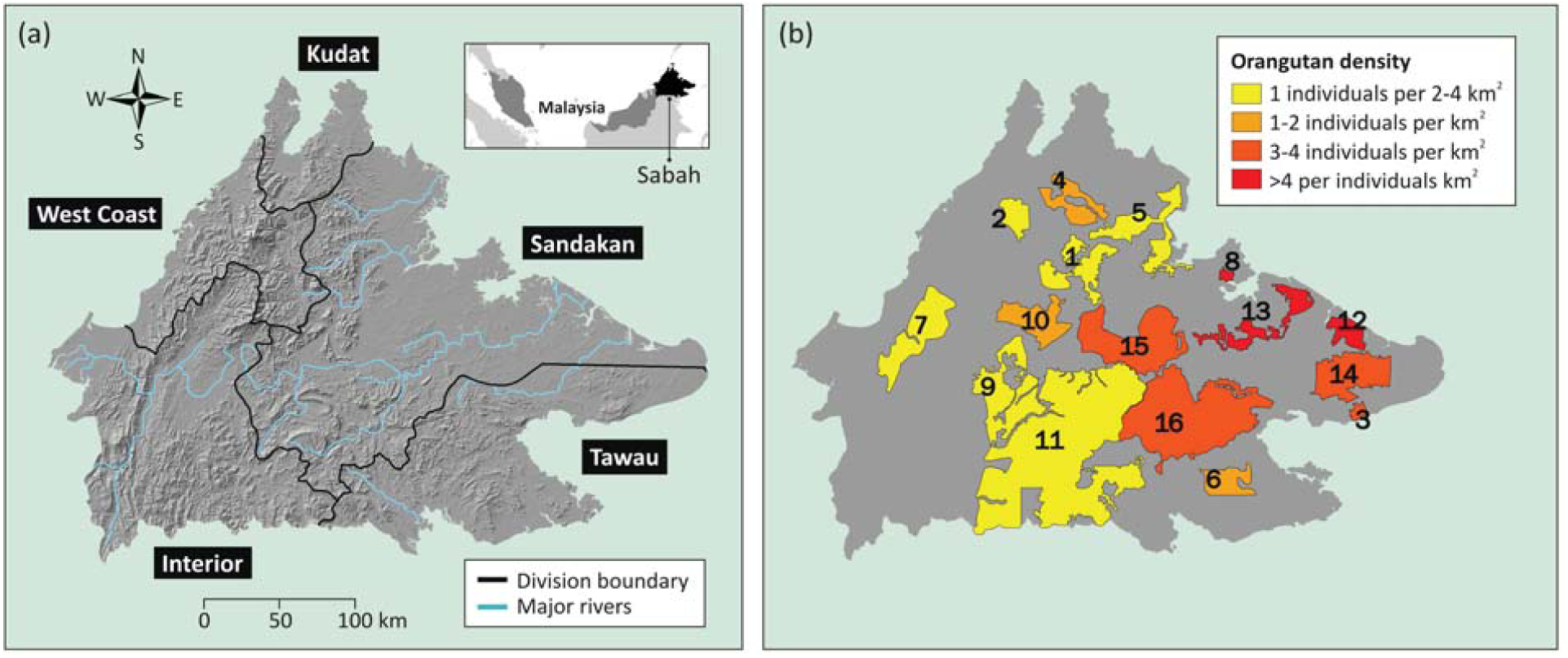
(a) Topological map of Sabah, Malaysia, with boundaries of administrative divisions and major rivers, and (b) 16 major orangutan populations identified by Ancrenaz *et al.* (2005), including: (1) Ulu Tungud, (2) Mount Kinabalu, (3) Silabukan, (4) Lingkabau, (5) Bongaya, (6) Ulu Kelumpang, (7) Crocker Range, (8) Sepilok, (9) Pinangah, (10) Trus Madi, (11) Kuamut, (12) Kulamba, (13) Kinabatangan, (14) Tabin, (15) Upper Kinabatangan, and (16) Segama.

## 2 METHODS

### 2.1 Study area

The study area covers the Malaysian state of Sabah (approximately 73,000 km^2^) (Figure 1a). In early 2000, 16 major (i.e. >50 individuals) orangutan populations throughout the state were identified by Ancrenaz *et al.* (2005) (Figure 1b). These populations were found either within fully protected areas (protected forest reserves (FR), Wildlife Sanctuaries (WS) or National Parks) or in logging concessions (commercial forest reserves (CFR)). The altitude of the study area ranged from 0 to 4000 meters, with low altitudes (0-100 meters) mainly occurring in the eastern part and higher altitudes (>500 meters) in the western part (Figure 1a). We divided the study area into grid cells with a spatial resolution of 1×1 km^2^.

### 2.2 Orangutan data

The present study utilized two types of orangutan data: nest counts and presence-absence data. The nest count data were obtained from line transect surveys (aerial and ground). The presence-absence data were derived from the line transect (aerial and ground) surveys and reconnaissance walks of nest observations.

The aerial surveys were conducted via helicopter during 1999-2002 and 2008-2012, with full details of the aerial survey methodology provided in Ancrenaz *et al*. (2005). The ground surveys were undertaken between 1999 and 2007 during several successive surveys and followed a standard established methodology to detect nests of great apes (Kühl 2007); these are extensively described by Ancrenaz *et al.* (2004b, 2005, 2010). The reconnaissance walk surveys were conducted sporadically between 1999 and 2012. For each survey method, we divided the data into three time periods: 1) pre-2003, 2) 2003-2007, and 3) 2008-2012, thus providing an analysis of the change in orangutan abundance every five years. This time interval conforms to the minimum inter-birth intervals (the time between consecutive offspring) of female Bornean orangutans (Knott *et al*. 2009). It also conforms to the time frames of orangutan conservation plans at a state level in Sabah (Sabah Wildlife Department 2011).

To facilitate the use of nest count data collected from various survey methods, we standardized the metric of orangutan nests across all surveys. For the ground transects surveys, we calculated the density of orangutan nests per km^2^ using the Distance sampling method, based on perpendicular distance of each nest to the transect (Buckland *et al*. 2005). For the aerial surveys, the data were mainly in the form of an aerial index value (*AI*) describing the number of nests detected per km flight. Following Ancrenaz *et al.* (2005), the density of orangutan nests per km^2^, i.e. *gnest*, was estimated via: log(*gnest*) = 4.7297 + 0.9796 log(*AI*). When all survey methods were combined there were approximately 4,500 grid cells or 1×1 km^2^ grids where orangutan nests had been observed. These data were then used to form variable *Y*_*i,j,t*_, where *Y*_*i,j,t*_ is a matrix arrays of survey periods (*t*), with each matrix consisting approximately 4500 rows of grid cells (*i*) and two columns of replicated nest counts from each survey (*j*).

To derive the occupancy of nests in each 1×1 km^2^ grid cell from line transect surveys (aerial and ground) and the reconnaissance walk surveys for each time period, we first divided the grid into sub-cells with a resolution of 200×200 m^2^. This is to avoid duplicated reports of the same clusters of nests (van Schaik *et al*. 2005). If the survey reported the occurrence of a nest within a sub-cell, we defined this as observed. If the survey reported the absence of a nest within a sub-cell or if there was no observation in that sub-cell, we defined this as unobserved. We then constructed a three dimensional matrix *Z*_*i,k,t*_ with three matrices of survey period (*t*), with each matrix consisting about 4,500 rows of grid cells (*i*) and 25 columns of observed and unobserved sub-cell (*k*).

### 2.3 Dynamic abundance model

#### 2.3.1 The model

We adapted the model developed by Dail & Madsen (2011) to analyze the orangutan population dynamics. Our model generalizes the negative binomial model for open populations and assumes that abundance patterns are determined by an initial territory establishment process followed by gains and losses resulting from births, mortality and dispersal. It also accounts for imperfect detection probability. Our model requires both spatial and temporal data and consists of three broad levels: (1) latent orangutan population, (2) latent orangutan nest, and (3) nest observation. The first level (latent orangutan population density) can be described as:

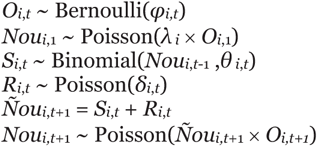

The second level (latent orangutan nest density) as:

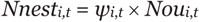

Finally, the third level (observed orangutan nest density and occupancy) as:

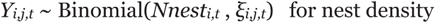

and *Znest*_*i,k,t*_ ∼ Bernoulli(*ρnest*_*i,k,t*_ × *Onest*_*i,t*_) for nest occupancy

where

*O*_*i,t*_ is the latent occurrence of orangutan at grid cell *i* in survey period *t*,

*Nou*_*i,t*_ is the latent number of orangutans at grid cell *i* in survey period *t*,

*S*_*i,t*_ is the latent number of survivors at grid cell *i* that do not emigrate between period *t* and *t*+1,

*R*_*i,t*_ is the latent number of recruits (including births and immigrants) at grid cell *i* between period *t* and *t*+1,

*Nnest*_*i,t*_ is the latent number of orangutan nests at grid cell *i* in survey period *t*,

*Onest*_*i,t*_ is the latent occupancy of orangutan nests at grid cell *i* in survey period *t*, derived as a binary value of *Nnest*_*i,t*_

*Y*_*i,j,t*_ is the observed nest count at grid cell *i* in survey period *t* from survey type *j*,

*Znest*_*i,k,t*_ is the observed nest occurrence at sub-grid cell *k* and grid cell *i* in survey period *t*

The parameters estimated from the model are the initial abundance rate at grid cell *i* (*λ*_*i*_), survival probability and recruitment rate at grid cell *i* between survey period *t* and *t*+1 (*θ*_*i,t*_ and *δ*_*i,t*_), the orangutan occupancy rate at grid cell *i* and survey period *t* (*φ*_*i,t*_), the scaling factor of the nest and the orangutan density at grid cell *i* and survey period *t* (*ψ*_*i,t*_), the probability of detecting orangutan nests from the line transects at grid cell *i* and survey period *t* for survey type *j* (*ξ*_*i,j,t*_, where *j*∈{aerial, ground}), and the probability of detecting orangutan nests from the line transects and reconnaissance walk surveys at sub-grid cell *k* and grid cell *i* and survey period *t* (*ρnest*_*i,k,t*_).

These parameters can be modeled by including grid cell-specific covariates. We modeled the initial abundance rate at grid cell *i* (*λ*_*i*_) as a function of altitude (*ALT*_*i*_), mean annual daily maximum temperature (*TEMP*_*i*,1_) and forest extent (*FOREST*_*i*,1_) in pre-2003, and the quadratic term for temperature, i.e.

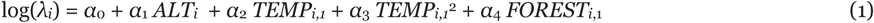

The occupancy rate at grid cell *i* in survey period *t* (*φ*_*i,t*_) and survival probability at grid cell *i* between period *t*-1 and *t* (*θ*_*i,t*_) were modeled in a similar manner as the initial abundance rate, i.e.

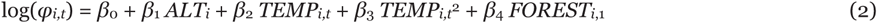

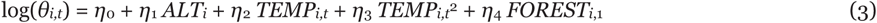

Altitude affects forest composition and structure, as well as fruit productivity, which in turn impacts abundance and survival of the Bornean orangutans (Wich *et al*. 2008). The extent of natural forest, i.e. primary old-growth forest and degraded forests that have not been clear cut, influences orangutan abundance since this species depends on tree cover to survive (Husson *et al*. 2009). Temperature affects the phenology of fruiting trees, which in turn affects orangutan abundance and survival rates (Hanya *et al*. 2013). We included the quadratic term for *TEMP* to test the preference of orangutan to occupy areas with intermediate values of mean annual maximum temperature. We did not include the temporal change in precipitation patterns as covariates because they are more difficult to discern than temperature, especially over short time intervals due to irregularities in the El Niño-Southern Oscillation (ENSO) cycle in this region (Malaysian Meteorological Department 2009). Description of the covariates used to explain the initial abundance, occupancy and survival rates are given in Table A1 (in Appendix A). Maps of the change in forest cover (*FOREST*) and mean annual daily maximum temperature (*TEMP*) for Sabah are shown in Figure A1.

The recruitment rate at grid cell *i* between period *t*-1 and *t* (*δ*_*i,t*_) was modeled as an autoregressive function of the number of individuals in grid cell *i* and the neighboring grid cells at the previous survey period (Besag 1974), i.e.

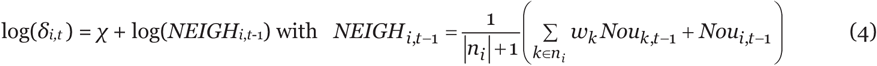

where *n*_*i*_ are the first-order neighbours surrounding grid cell *i* and *w*_*k*_ is a binary indicator (1 or 0) of whether cell *i* is connected to cell *k*∈*n*_*j*_. The binary indicator *w*_*k*_ was introduced to take into account the effect of large rivers on orangutan dispersal. Large rivers influence the population genetic structure of orangutans since orangutans cannot swim or cross large water bodies (Jalil *et al*. 2008). A study around the Kinabatangan River (Jalil *et al*. 2009) found that orangutan genetic samples from each side of the river were significantly differentiated by high molecular variance. We used a spatial map of major rivers within the study area (Figure 1a) to determine barriers to orangutan dispersal. To build *w*_*k*_, we first constructed a vector of straight lines that connect the centre point of cell *i* and the centre point of each adjacent cell *k*∈*n*_*j*_ (Santika *et al*. 2015). This is to simulate the possible dispersal routes taken by an orangutan from cell *i* to the surrounding cells. We then intersected this line with the river barrier layer. We assumed *w*_*k*_=0 if at least one intersection was found within cell *k*∈*n*_*j*_ (i.e. rivers prevent orangutan dispersal from cell *i* to cell *k*) and *w*_*k*_=1 if no intersection was found.

In earlier studies, the density of orangutans at grid cell *i*, i.e. *gou*_*i*_, has typically been estimated by the following equation

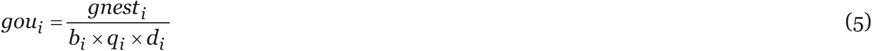

where *b*_*i*_ is the proportion of nest builders, i.e. juveniles less than around 3 years of age are unlikely to build nests and share nests with their mothers (Prasetyo *et al*. 2009), *q*_*i*_ is the daily rate of nest production, and *d*_*i*_ is the nest decay rate or the number of days a nest remains visible. Based on previous studies in Sabah, the proportion of nest builders has been estimated as 0.9 (Ancrenaz *et al*. 2005, 2010). The average daily rate of nest production for Sabah has been estimated as 1.01 (Ancrenaz *et al*. 2005), but this can fluctuate depending on the level of forest disturbance, i.e. between primary and logged over forest (Rayadin & Saitoh 2009). Generally, the multiplication of *b*_*i*_ and *q*_*i*_ results in a value around 1. The nest decay rate is much more uncertain, however, ranging between around 85 to 850 days (Mathewson *et al*. 2008; Marshall & Meijaard 2009) and has been shown to vary across different forest types, altitude and climate (Ancrenaz *et al*. 2004a, b). Hence, to take into account the variability in the total denominator of Eq. (5) across different grid cells *i* and survey periods *t*, we modeled *ψ*_*i,t*_ as

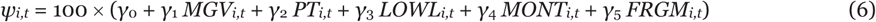

where *MGV*_*i,t*_ is a binary variable denoting whether or not the majority of forest at grid cell *i* and time *t* are mangrove forest, and similarly *PT*_*i,t*_ for peat forest, *LOWL*_*i,t*_ for lowland forest (altitude <500 m), *MONT*_*i,t*_ for montane forest (altitude ≥500 m), and *FRGM*_*i,t*_ for highly fragmented forest (<25 ha per km^2^).

The probability of detecting orangutan nests at grid cell *i* and time *t* and for survey *j* (*j*∈{aerial, ground}), i.e. *ξ*_*i,j,t*_, was modeled constant for each survey type, such that

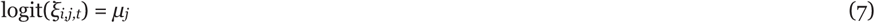

Finally, the probability of detecting orangutan nests at sub-grid cell *k* and grid cell *i* and time *t* for line transects and other targeted surveys, i.e. *ρnest*_*i,k,t*_, was modeled constant, such that

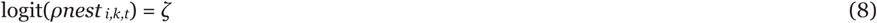

Both parameters *μ*_*j*_ and *ζ* are to be estimated.

#### 2.3.2 Model fitting and evaluation

We used WinBUGS Version 1.4.3 (Lunn *et al*. 2000) to estimate the parameter posterior distributions and the regression coefficients for *λ*_*i*_, *φ*_*i,t*_, *θ*_*i,t*_, *δ*_*i,t*_, *ψ*_*i,t*_, *ξ*_*i,j,t*_, and *ρnest*_*i,k,t*_. The WinBUGS code for the dynamic abundance model is provided in Appendix B. We assumed a uniform prior for each variable explaining the initial abundance rate, occupancy rate, and survival probability (U[-4,4] for parameters in Eq. (1-3)), recruitment rate (U[-4,4] for parameter in Eq. (4)), the scaling factor for orangutan and nest abundance (U[-10,10] for parameters in Eq. (6)), and detection probability (U[-4,4] for parameters in Eq. (7-8)), as described in Appendix B.

We ran three Markov chain Monte Carlo (MCMC) chains, where each chain consists of 100,000 iterations and the first 50,000 were discarded as burn-in. To improve convergence and to reduce the autocorrelation in the MCMC chain, we standardized all variables prior to model fitting. Prior to fitting the model to the data, we tested the correlation among the environmental variables explaining *λ, φ*_*t*_ and *θ*_*t*_, i.e. variables *ALT, TEMP* and *FOREST.* Convergence for each model parameter was assessed from the values of Rhat statistics and visualization of the chain plot of the MCMC iterations. Rhat values around 1 and the absence of seasonality within each chain plot and overlap among the chains indicate convergence.

We assessed the predictive performance of the dynamic abundance model by comparing the actual nest density at the final time period (2008-2012) with the simulated nest predictions obtained by fitting the model to the first two time periods of the data (pre-2003 and 2003-2007). For each simulated prediction, we calculated the Pearson’s correlation coefficient *r* and also fitted a linear regression between the predicted values and the observed values to calculate the *R*^2^ value (Bahn & McGill 2013).

### 2.4 Orangutan abundance and land use change

We assessed how the orangutan abundance (obtained from the simulated predictions) changes with the change in land uses and considered four land use categories: (a) protected areas (PA) (which includes protection forest reserves under the Sabah Forestry Department (class I based on the Malaysian Permanent Forest Reserves classifications), mangrove forest reserves (class V), virgin jungle reserves (class VI) and wildlife reserves (class VII); wildlife sanctuaries under the Sabah Wildlife Department and national parks under Sabah Parks), (b) commercial forest reserves (CFR) designated for sustainable logging (class II), (c) industrial timber plantation concessions (ITP), (d) oil palm plantations (OPP), and (e) area outside PA, CFR, ITP and OPP (Other) (Figure A2). We obtained spatial boundary data for PA, CFR and ITP for 2000, 2006 and 2012 from the Sabah Forestry Department Annual Reports, and spatial data for OPP from the Malaysian Palm Oil Board (MPOB).

## 3 RESULTS

### 3.1 Model predictive performance and surveys’ detection rates

The model performed relatively well with a moderate correspondence between the simulated nest predictions for the final time period (2008-2012) and the actual observations for this time period. The average Pearson correlation coefficient is *r*=0.682 and the average *R*^2^ is 0.604.

Aerial surveys had a higher probability of detecting orangutan nests per km^2^ (75.6%, 95% credible interval (CI): 70.9%-80.4%) compared to line transect ground surveys (47.5%, 95% CI: 41.1%-53.8%) (Table 1). This is consistent with the fact that the aerial surveys were conducted in areas where orangutans were known to be present, given the cost for operating helicopters. In contrast, the locations of ground transect were selected more randomly in locations with unknown orangutan abundance or presence.

**Table 1.** Posterior means and the 95% credible interval of the mean for each parameter of the orangutan dynamic abundance model.

| Sub-model | Scale | Variable (Parameter) | Posterior mean | 95% credible interval of mean |
| --- | --- | --- | --- | --- |
| Initial abundance in pre-2003 ($\lambda_i$ in Eq. (1)) | Log | Intercept ($\alpha_0$) | 1.141 | (0.821, 1.562) |
| | | *ALT* ($\alpha_1$) | -1.057 | (-2.126, -0.592) |
| | | *TEMP* ($\alpha_2$) | 0.812 | (0.415, 1.065) |
| | | *TEMP*<sup>2</sup> ($\alpha_3$) | -0.564 | (-0.872, -0.271) |
| | | *FOREST* ($\alpha_4$) | 0.306 | (0.067, 0.686) |
| Occupancy rate every five years ($\varphi_{i,t}$ in Eq. (2)) | Logit | Intercept ($\beta_0$) | 1.112 | (0.772, 1.517) |
| | | *ALT* ($\beta_1$) | -1.121 | (-2.189, -0.762) |
| | | *TEMP* ($\beta_2$) | 0.719 | (0.351, 1.214) |
| | | *TEMP*<sup>2</sup> ($\beta_3$) | -0.532 | (-0.952, -0.190) |
| | | *FOREST* ($\beta_4$) | 0.431 | (0.125, 0.767) |
| Survival probability every five years ($\theta_{i,t}$ in Eq. (3)) | Logit | Intercept ($\eta_0$) | 1.762 | (1.472, 2.162) |
| | | *ALT* ($\eta_1$) | -0.012 | (-0.158, 0.026) |
| | | *TEMP* ($\eta_2$) | 0.365 | (0.024, 0.683) |
| | | *TEMP*<sup>2</sup> ($\eta_3$) | -0.201 | (-0.520, -0.062) |
| | | *FOREST* ($\eta_4$) | 0.714 | (0.424, 1.085) |
| Immigration rate every five years ($\delta_{i,t}$ in Eq. (4)) | Log | Intercept ($\chi$) | -1.931 | (-2.325, -1.527) |
| Scaling factor of the nest and the orangutan density ($\psi_{i,t}$ in Eq. (6)) | Normal | Intercept ($\gamma_0$) | 2.036 | (1.572, 2.452) |
| | | *MGV* ($\gamma_1$) | 0.204 | (0.052, 0.426) |
| | | *PT* ($\gamma_2$) | 0.114 | (0.008, 0.187) |
| | | *LOWL* ($\gamma_3$) | 0.035 | (-0.013, 0.098) |
| | | *MONT* ($\gamma_4$) | -0.134 | (-0.348, -0.078) |
| | | *FRGM* ($\gamma_5$) | 0.092 | (-0.017, 0.132) |
| Nest detection rate from line transect surveys (density) ($\xi_{i,j,t}$ in Eq. (7)) | Logit | Intercept ($\mu_{\text{aerial}}$) | 1.131 | (0.895, 1.412) |
| | | Intercept ($\mu_{\text{ground}}$) | -0.102 | (-0.361, 0.158) |
| Nest detection rate from line transect and targeted surveys (occurrence) ($\rho_{nest,i,k,t}$ in Eq. (8)) | Logit | Intercept ($\zeta$) | -1.289 | (-1.712, -0.942) |

### 3.2 Orangutan density estimate

The coefficients for the climate and environmental predictors in the model of initial abundance (in pre-2003) provide an indication of the magnitude of their effects on the long-term orangutan abundance rates (Table 1). The model indicates that orangutan abundance per km^2^ prior to 2003 correlates strongly with altitude and mean annual daily maximum temperature. Orangutans are most abundant in lowland areas with moderate temperature. Altitude and mean annual daily maximum temperature also had large effects on orangutan occupancy rates in each five year period. The effects of the climate and environmental predictors on survival rates per km^2^ for every five year period, however, were quite different from the initial abundance and occupancy rate. Natural forest cover had the largest effect, with survival rates highest in areas with extensive forest cover. The probability of orangutans migrating more than 1 km distance every five years was generally low.

Our model predicted that prior to 2003, orangutans were abundant mainly in the eastern coastal and the central parts of Sabah (Figure 2). Our model indicated that the total number of orangutans has declined from about 14,000 individuals to about 10,000 between 2003 and 2012, equating to a 27.8% reduction in their numbers over the last ten years (Table 2).

**Table 2.**
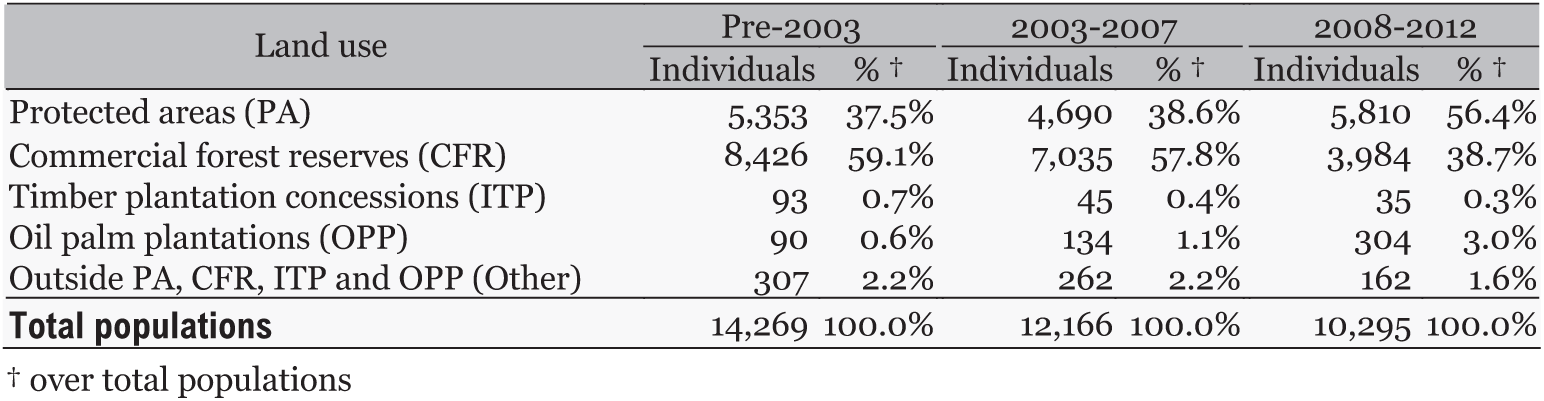
The estimated number of individuals within the study area across different land-use categories.

**Figure 2.**
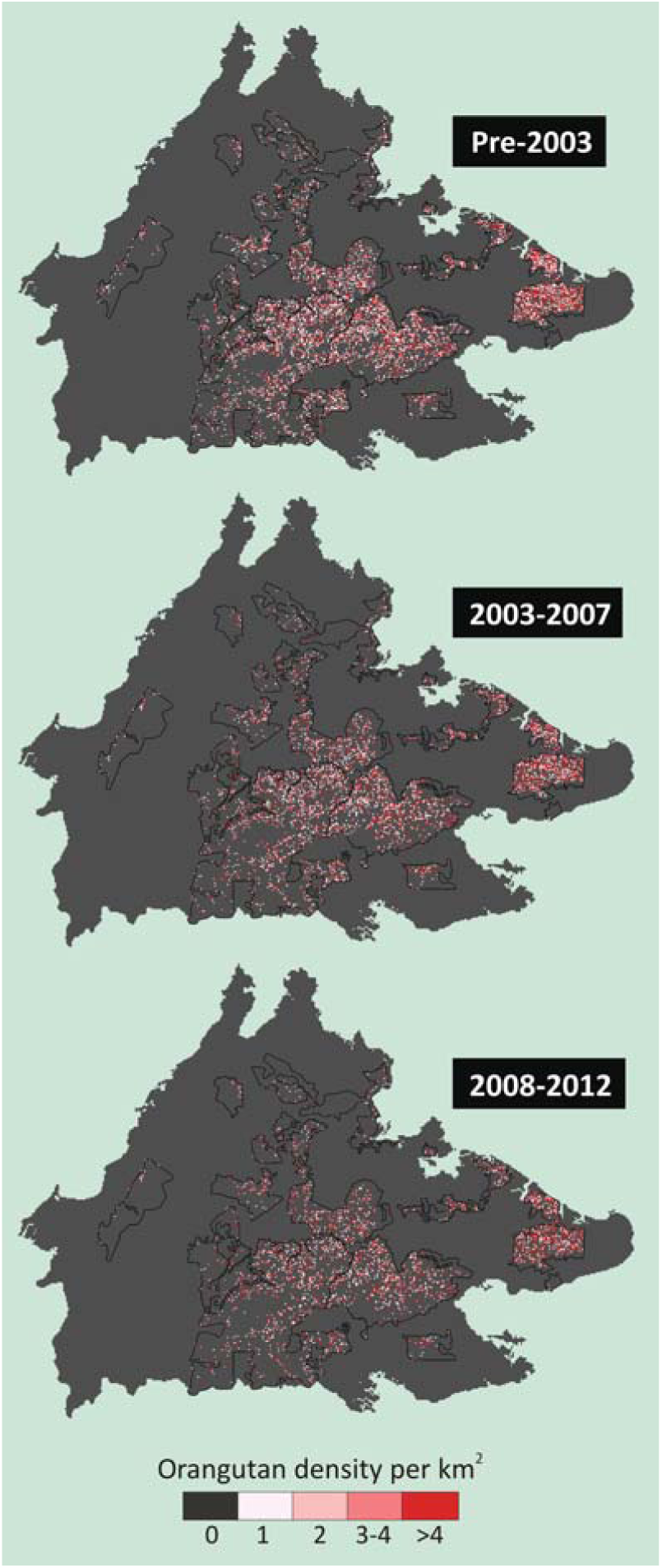
The change in orangutan density per km^2^ in Sabah from three time periods during 1999-2012. Black lines indicate the boundaries of the 16 populations identified by Ancrenaz *et al.* (2005).

The rates of decline in the total number of individuals varied for the different populations. Populations 3, 5, 9 and 11 had the highest declining rates (≥75th percentile or ≥15% every five years), whereas populations 12 and 14 had the lowest rates (<25th percentile or <10% every five years) (Figure 3a and Table A2). High declining rates appeared to be attributed mainly to extensive habitat loss (population 5, 9 and 11), followed by small population size and a high level of isolation (population 3) (Figure 3b).

**Figure 3.**
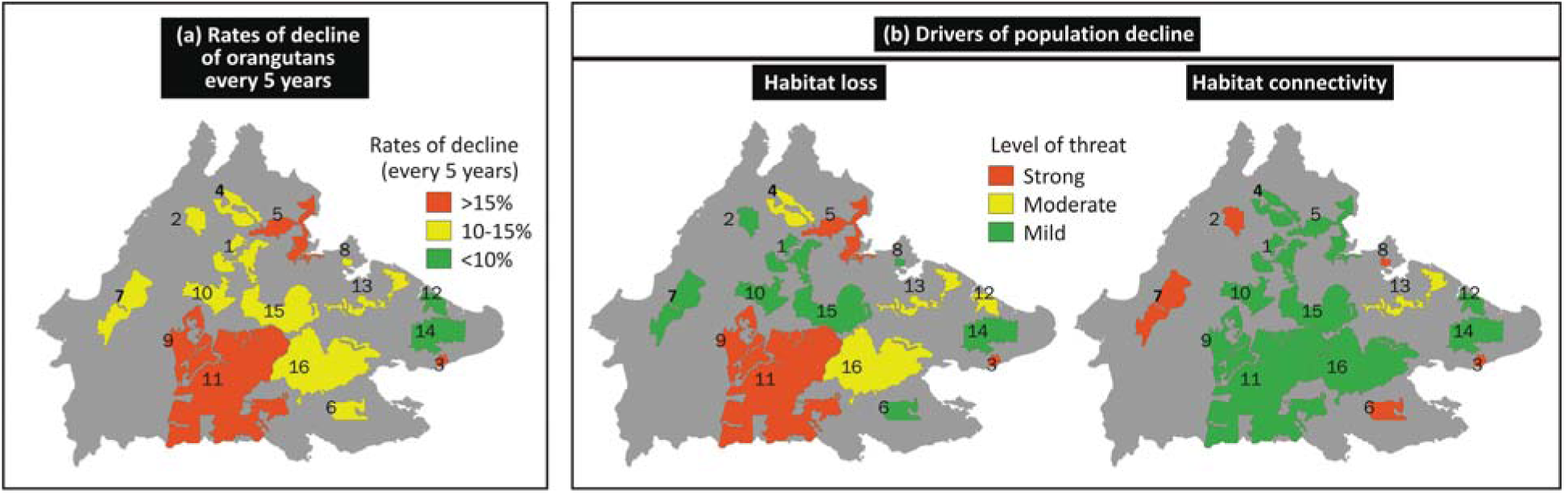
(a) Estimated mean declining rates in orangutan numbers every five years within each of 16 populations identified by Ancrenaz *et al.* (2005), grouped according to percentile range: ≥75th percentile (≥15%), 25-75th percentile (10-15%), and <25th percentile (<10%). (b) Drivers of population decline, which include habitat loss and habitat connectivity. Habitat loss was defined as the loss of natural forest (ha per km2 every five years) within the area covered by each population. Habitat connectivity was defined as the interaction between the reciprocal size of habitat patch where the population lives and distance to nearest population habitats, where large value indicates isolated population. Different levels of threats were determined based on percentile range, i.e. Strong: >75th percentile, Moderate: 25-75th percentile, and Mild: <25th percentile.

Our data indicated that prior to 2003, 59% of the total orangutan population in Sabah resided inside commercial forest reserves (CFR) and 37% were inside protected areas (PA) (Table 2). As the extent of protected areas has increased recently, the populations inside PA now make up about 56% of the orangutan population found within the study area, while 38% remain inside the CFRs. The establishment of oil palm plantations (OPP) has occurred in several areas occupied by orangutan, accounting for less than 1% of the total orangutan populations prior to 2003, but then increasing to 3% in 2008-2012.

### 3.3 Scaling factor of orangutan and nest density

The intercept of the scaling factor that describes the relationships between the number of orangutans and the number of nests (*γ*_0_) was estimated to be 2.036 (Table 1). Assuming the value of the proportion of nest builders and the daily rate of nest production is close to 1, this implies that the nest decay rate across different grid cells within 1×1 km^2^ grids within the study area was about 204 days. This conforms to the value of 202 days estimated by Ancrenaz *et al.* (2005) for the Kinabatangan area. The scaling factors varied, although not substantially, across different forest types. Nests within mangrove and swamp forest appeared to have the longest time to decay (220 days on average), followed respectively by mountain forest (217 days) and lowland forest (207 days). Nests in highly fragmented forest generally had a decay rate of about 213 days.

## 4 DISCUSSION

For many highly threatened species, conservation plans are usually made at national or regional level, and to identify suitable strategies for their monitoring, management and recovery requires understanding of the change in their abundance over the extent of interest and landscape drivers of this change. However, data on species abundance obtained from repeated surveys over large areas are often lacking due to limited resources (Field *et al*. 2005). Instead, time series abundance data on a species are typically available locally. The utility of such data has so far been limited either (1) to generate a local population trend by ensuring consistent data and survey methodologies through time (Forcada *et al*. 2006), i.e. a ‘dynamic but local’ study, or (2) to generate a static species abundance model, i.e. models that estimate species abundance based on current environmental predictors, over a wide area allowing for the use of various sources of data to represent the species current abundance distribution, i.e. a ‘broad but statić study (Guralnick *et al*. 2005). The study of changes in species abundance over large areas, i.e. ‘broad and dynamić, are rare due to the lack of suitable methodologies for analyzing data commonly available. Our analysis is a significant advance over current approaches by allowing a complete utilization of different time series datasets on species abundance across different localities whilst accounting for varying error rates in detection probability inherent to each dataset. It provides a reliable measure as well as an explanation of species persistence through time, the aspect of most concern in conservation of a species.

In this study we demonstrate how different survey protocols yielded a different probability of detecting orangutan, with aerial surveys having the highest detection probability, followed by line transects ground surveys. The importance of accounting for detection error, especially related to survey effort, has been highlighted in previous literature (Kéry *et al.* 2005; Chen *et al.* 2009). Our study corroborates previous works on the need of accounting for detection error in modelling species distribution.

Our study found that the primary orangutan population in Sabah has been declining by more than 25% over the last ten years. This is higher than the previous estimate of a 35% decline over two decades suggested by Ancrenaz *et al.* (2005), in which the authors compared the estimated populations in 1999-2002 with an earlier estimate from the 1980s by Payne (1987). A genetic study by Goossens *et al.* (2006) estimate a 95% decline in the last 200 years for populations in the Lower Kinabatangan, with population collapses mostly occurring in recent decades. Our model also predicted that prior to 2003 about 14,000 individuals existed over the entire state of Sabah, and this is consistent with Ancrenaz *et al*. (2005) who estimated between 8,000 and 18,000 individuals in early 2000. Our estimate for population 8 (Sepilok FR), however, was lower (2.13 individuals per km^2^) than the estimate made by the authors (5 individuals per km^2^) (Table A2). This is mainly because unlike the previous study our analysis does not take into account the fact that Sepilok FR has been an orangutan site in which rehabilitated orangutans are released. Sepilok Rehabilitation Center has received more than 600 orangutans over the past fifty years (Kuze *et al.* 2008) and about of third of them was released in a 40 km^2^ forested area during this time (Fernando 2001; Russon 2009), although post-release mortality is unknown.

The loss of natural forest, i.e. both undisturbed primary old-growth forest and degraded forests exploited for timber, had the largest impact on orangutan survival rates, followed by rising mean annual daily maximum temperature. The proportion of orangutan populations occurring in areas where oil palm plantation were later established has increased over the last ten years due to the expansion of this monoculture crop, while about a third of these populations currently remain inside the CFR designed for sustainable logging. Recent studies have shown that forests that are heavily logged or subjected to fast conversion cannot maintain orangutan populations (Husson *et al*. 2009; Ancrenaz *et al*. 2010). Thus, as the expansion of oil palm and pulp and paper industries is expected to continue in the coming decades in response to global demand for food, fiber and fuel (Koh & Wilcove 2008), without proper land use planning that accounts explicitly for orangutan habitats, it is likely that this development will result in further declines of orangutan populations. Additionally, the situation may worsen as climate conditions are predicted to be more extreme due to both global and regional climate change (Malaysian Meteorological Department 2009; Struebig *et al*. 2015).

Our findings highlight the importance of natural forest for long-term orangutan survival. Maintaining ecological connectivity between forest patches is also important as our study shows that the orangutan migration rates across more than 1 km distance is generally low. Increased terrestrial movement in a human-made vegetation matrix may also increase conflict with people and exposure to new diseases (Loken *et al*. 2013; Ancrenaz *et al*. 2014). The Sabah government’s recent decision to expand their protected area extent from 16.0% of their land mass to 22.4% (mainly by connecting Maliau Basin FR and Danum Valley FR) has resulted in the protection of 60% of the orangutan populations within the study area. As the government plans to further expand their protected areas network to 30% of the land mass over the next ten years (WWF Malaysia 2014), our study can be used to guide the assessment of suitable areas for the expansion.

In this study we noted some key limitations that present future challenges. We used forest type as a predictor of the variability of nest decay rates while studies have shown that the rate can also vary with nest height and annual rainfall (Mathewson *et al*. 2008). If detailed data on nest height had been available, it would still not be possible to generalize this information over the grid scale used in this study (1×1 km^2^). Furthermore, including mean annual rainfall to explain both the nest decay rates (at nest observation level) and the initial population abundance and survival rates (at the orangutan population level), will likely lead to a confounding measures and unreliable parameter estimates, unless the range of prior distributions for the parameters are known with certainty. Additionally, some of the orangutan surveys were conducted only once in a site within a time period, and this can lead to a less reliable probability of detection estimates. Thus, our study would benefit from alterations to survey designs to facilitate more robust estimation of detectability.

Our analysis can be improved by including a sex-specific dispersal rate. A study from Lower Kinabatangan in early 2000 indicates that the orangutan populations were above the carrying capacity of the habitat with the number of males substantially greater than the number of females, most likely due to difference in male versus female dispersal patterns (Marshall *et al.* 2009). A genetic analysis from seven Borneo populations suggests that male dispersal distances are several-times higher than those of females, leading to strong male-biased dispersal in orangutans (Nietlisbach *et al*. 2012). Sex-biased dispersal has recently been incorporated into dynamic metapopulation models, i.e. models that are based on presence-absence data (Dolrenry *et al*. 2014), and the incorporation of sex-biased dispersal in the dynamic abundance modeling presents a future research challenge.

With the number of species listed in the IUCN Red List having increased over the years (Hoffmann *et al*. 2010), there is growing pressure for conducting evidence-based assessments on species persistence to inform policy at regional and national levels (Hutchings *et al*. 2012). As the availability of species survey data continues to improve (e.g., through emerging citizen science projects coupled with online species databases (Silvertown 2009)) coupled with improved data sharing between scientists, and governmental and non-governmental agencies (Tenopir *et al*. 2011), our methodology provides a valuable tool for analyzing diverse data sources in an integrative manner.

## ACKNOWLEDGEMENTS

This study is part of the Borneo Futures initiative and was supported by the Australian Research Council (Centre of Excellence and Future Fellowship programs) and the Arcus Foundation.

