## Appendix for "The power of mixed survey methodologies for detecting decline of the Bornean orangutan"

**Table A1.** Description of the covariates used to explain the initial abundance, occupancy and survival rates in the dynamic population model. All data were adjusted to a spatial resolution of 1×1 km².

| Covariate | Static/ Dynamic | Scale | Data preparation and sources |
|---|---|---|---|
| Altitude (*ALT*) | Static | Meters | Shuttle Radar Topography Mission (SRTM) digital elevation model (Jarvis *et al.* 2008) |
| Natural forest extent (*FOR*) | Dynamic (pre-2003, 2003-2007, 2008-2012) | Hectares | The extent of natural forest map for 2000 was obtained from the Sabah Forestry Department (2011) and was used to represent the natural forest extent pre-2003. The extent of natural forest for the subsequent time periods were derived by overlaying the extent of natural forest pre 2003 and the annual forest cover loss between 2000 and 2012 derived from the Global Forest Change data (Hansen *et al.* 2013). The annual forest loss data has a spatial resolution of 30×30 m² and comprises binary values, where "1" denotes clearing of forest within a 30×30 m² grid and "0" otherwise. |
| Mean annual daily maximum temperature (*TEMP*) | Dynamic (pre-2003, 2003-2007, 2008-2012) | °C | The mean annual daily maximum temperature for pre-2003, 2003-2007, and 2008-2012 were obtained by applying a thin plate smoothing spline over daily maximum temperature data recorded at 180 meteorological stations and agriculture offices across Borneo (BPS 2014; NOAA 2014) and taking into account the geographical position and altitude of each station. We used the ANUSPLIN software package to run the spline smoothing (Hutchinson 2004; Hijmans *et al.* 2005). |

**Table A2.** The change in the density of orangutan per km² within each of the ten populations identified by Ancrenaz *et al.* (2005). Rows highlighted in dark grey, light grey and yellow represent populations with mean declining rates every five years of ≥15% (≥75th percentile), 10-15% (25-75th percentile) and <10% (<25th percentile), respectively.

| Population | Orangutan density per km² | | | | Mean declining rates every five years |
|---|---|---|---|---|---|
| | Early 2000 (MA) | Pre-2003 | 2003-2007 | 2008-2012 | |
| 1 | 0.04 | 0.08 | 0.07 | 0.06 | 13.4% |
| 2 | 0.25 | 0.17 | 0.15 | 0.13 | 12.5% |
| 3 | 0.58 | 0.38 | 0.32 | 0.27 | 15.7% |
| 4 | 0.33 | 0.22 | 0.18 | 0.16 | 14.6% |
| 5 | 0.19 | 0.18 | 0.15 | 0.13 | 16.7% |
| 6 | 0.30 | 0.19 | 0.16 | 0.14 | 14.1% |
| 7 | 0.2 | 0.16 | 0.14 | 0.12 | 13.4% |
| 8 | 5.00 | 2.13 | 1.86 | 1.66 | 11.7% |
| 9 | 0.22 | 0.19 | 0.16 | 0.13 | 17.3% |
| 10 | 0.41 | 0.38 | 0.31 | 0.28 | 14.0% |
| 11 | 0.06 | 0.39 | 0.34 | 0.27 | 16.7% |
| 12 | 2.94 | 3.90 | 3.64 | 3.31 | 7.9% |
| 13 | 2.74 | 2.79 | 2.47 | 2.07 | 13.8% |
| 14 | 1.26 | 1.37 | 1.26 | 1.15 | 8.4% |
| 15 | 1.03 | 1.15 | 1.04 | 0.93 | 10.1% |
| 16 | 0.99 | 0.67 | 0.60 | 0.52 | 11.9% |

##### References Table A1-2

Ancrenaz, M., Gimenez, O., Ambu, L., Ancrenaz, K., Andau, P. *et al.* (2005) Aerial surveys give new estimates for orangutans in Sabah, Malaysia. *PLoS Biology* **3**, e3.

Bureau of Statistics Indonesia (BPS) (2014) *Kabupaten Dalam Angka*. Jakarta.

Hansen, M.C., Potapov, P.V., Moore, R., Hancher, M., Turubanova, S.A. *et al.* (2013) High-resolution global maps of 21st-century forest cover change. *Science* **342**, 850-853.

Hijmans, R.J., Cameron, S.E., Parra, J.L., Jones, P.G. & Jarvis, A. (2005) Very high resolution interpolated climate surfaces for global land areas. *International Journal of Climatology* **25**, 1965-1978.

Hutchinson, M. (2004) *ANUSPLIN Version 4.3*. The Australian National University, Canberra.

Jarvis, A., Reuter, H.I., Nelson, A. & Guevara, E. (2008) *Hole-filled SRTM for the Globe Version 4*. Available from https://srtm.csi.cgiar.org. Accessed 10 April 2018.

National Oceanic and Atmospheric Administrations USA (NOAA) (2014) *Climate Data Online (CDO)*. Available from https://www.ncdc.noaa.gov/cdo-web. Accessed 4 March 2018.

Sabah Forestry Department (2011) *Fact Sheets of Forest Reserves in Sabah*. Sandakan, Sabah.

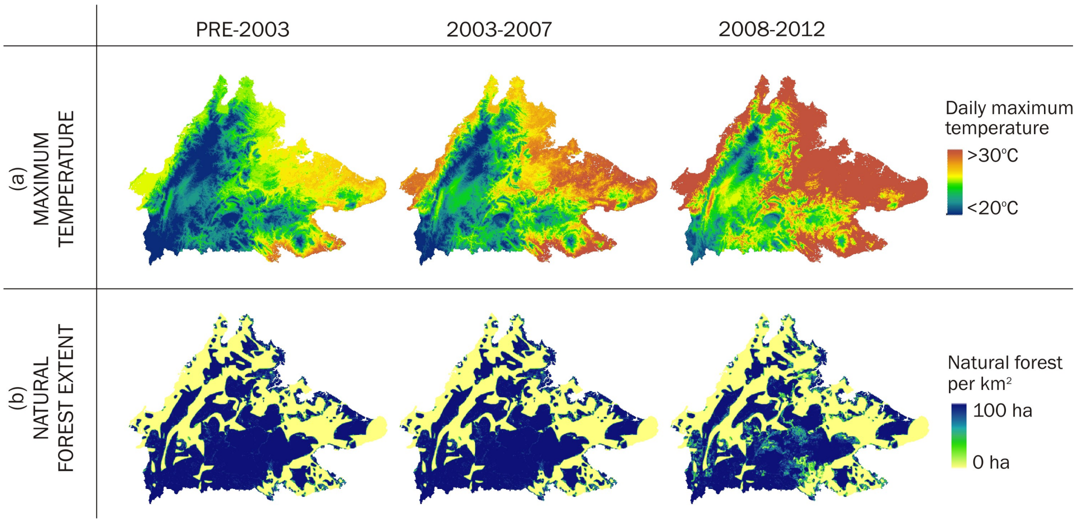


**Fig. A1.** The change in (a) natural forest extent per km² and (b) the mean annual daily maximum temperature, in Sabah over the last decade.

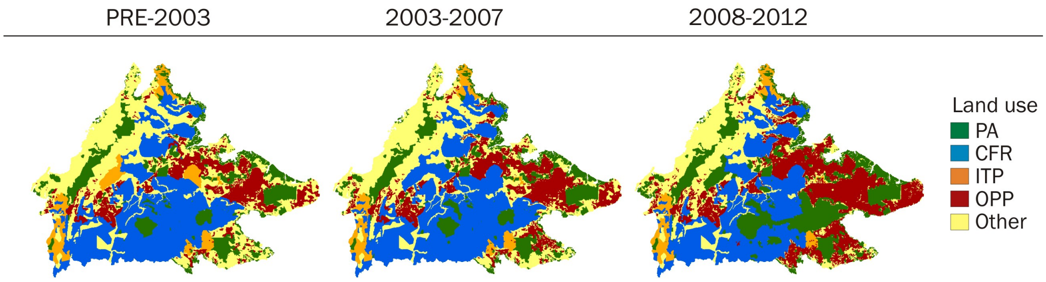


**Fig. A2.** The change in land use allocations for protected areas (PA), commercial forest reserves (CFR), timber plantation concessions (ITP), and oil palm plantations (OPP), and Other (outside PA, CFR, ITP and OPP), over the last decade.

### Appendix B. WinBUGS Code

```
#--------------------------------------------------------------------------------
# The power of mixed survey methodologies for detecting decline of the Bornean
# orangutan

# Authors: Truly Santika, Kerrie A. Wilson, Erik Meijaard & Marc Ancrenaz
#--------------------------------------------------------------------------------


# WINBUGS code (core component)

PopModel = "model
{
# PRIORS
for (i in 1:5)
       {
       alpha[i] ~ dunif(-4,4)
       beta[i] ~ dunif(-4,4)
       eta[i] ~ dunif(-4,4)
       }

gamma[1] ~ dunif(0,10)
for (i in 2:6)
       {
       gamma[i] ~ dunif(-10,10)
       }

for (i in 1:2)
       {
       mu[i] ~ dunif(-4,4)
       }

zeta ~ dunif(-4,4)
chi ~ dunif(-4,4)


# INITIAL ABUNDANCE (TIME PERIOD 1)
for (i in 1:G)
       {
       # LEVEL 1 (LATENT ORANGUTAN POPULATION)

       # Latent abundance and distribution
       log(lambda[i,1]) <- alpha[1]+alpha[2]*ALT[i]+alpha[3]*TEMP[i,1]+alpha[4]*TEMP2[i,1]
       +alpha[5]*FOREST[i,1]

       logit(phi[i,1]) <- beta[1]+beta[2]*ALT[i]+beta[3]*TEMP[i,1+beta[4]*TEMP2[i,1]
       +beta[5]*FOREST[i,1]

       O[i,1] ~ dbern(phi[i,1])
       Nu[i,1] <- lambda[i,1]*O[i,1]
       N.ou[i,1] ~ dpois(Nu[i,1])

       # LEVEL 2 (LATENT ORANGUTAN NEST POPULATION)

       # Abundance and nest relationship
       psi[i,1] <- gamma[1]+gamma[2]*MGV[i,1]+gamma[3]*PT[i,1]+gamma[4]*LOWL[i,1]
       +gamma[5]*MONT[i,1]+gamma[6]*FRGM[i,1]

       N.nest[i,1] ~ dpois(N.ou[i,1]*psi[i,1]*100)
       O.nest[i,1] <- step(N.nest[i,1]-1)

       # LEVEL 3 (NEST OBSERVATION)

       # Count data for (1) aerial surveys and (2) ground line transects
       for (j in 1:2)
              {
              logit(xi[i,j,1]) <- mu[j]
              Y[i,j,1] ~ dbin(xi[i,j,1],N.nest[i,1])  # Count data
              }
```

```
        # Observed/unobserved data for aerial and ground transect surveys and reconnaissance
        # walks

        for (k in 1:K)
                {
                logit(rhonest[i,k,1]) <- zeta
                zi[i,k,1] <- O.nest[i,1]*rhonest[i,k,1]
                Z.nest[i,k,1] ~ dbern(zi[i,k,1])  # Occurrence data
                }
        }

# SUBSEQUENT ABUNDANCE (TIME PERIOD 2 AND 3)
for (i in 1:G)
        {
        for (t in 2:T)
                {
                # LEVEL 1 (LATENT ORANGUTAN POPULATION)

                # Colonization rate
                nbr[i,1,t-1] <- 0
                for (j in 1:TOTNEIGH[i])
                        {
                        nbr[i,j+1,t-1] <- nbr[i,j,t-1]+N.ou[NEIGHBOUR[i,j],t-1]
                        }
                neigh[i,t-1] <- nbr[i,(TOTNEIGH[i]+1),t-1]/TOTNEIGH[i]

                log(delta[i,t-1]) <- chi+log(neigh[i,t-1])
                R[i,t] ~ dpois(delta[i,t-1])

                # Survival rate
                logit(theta[i,t]) <- eta[1]+eta[2]*ALT[i]+eta[3]*TEMP[i,t]+eta[4]*TEMP2[i,t]
                +eta[5]*FOREST[i,t]

                S[i,t] ~ dbin(theta[i,t], N[i,t-1])
                lambda[i,t] <- S[i,t] + R[i,t]

                # Occupancy
                logit(phi[i,t]) <- beta[1]+beta[2]*ALT[i]+beta[3]*TEMP[i,t]
                +beta[4]*TEMP2[i,t]+beta[5]*FOREST[i,t]

                O[i,t] ~ dbern(phi[i,t])

                # Latent abundance
                Nu[i,t] <- lambda[i,t]*O[i,t]
                N.ou[i,t] ~ dpois(Nu[i,t])

                # LEVEL 2 (LATENT ORANGUTAN NEST POPULATION)

                # Abundance and nest relationship
                psi[i,t] <- gamma[1]+gamma[2]*MGV[i,t]+gamma[3]*PT[i,t]+gamma[4]*LOWL[i,t]
                +gamma[5]*MONT[i,t]+gamma[6]*FRGM[i,t]

                N.nest[i,t] ~ dpois(N.ou[i,t]*psi[i,t]*100)
                O.nest[i,t] <- step(N.nest[i,t]-1)

                # LEVEL 3 (NEST OBSERVATION)

                # Count data for (1) aerial surveys and (2) ground line transects
                for (j in 1:2)
                        {
                        logit(xi[i,j,t]) <- mu[j]
                        Y[i,j,t] ~ dbin(xi[i,j,t],N.nest[i,t])  # Count data
                        }

                # Observed/unobserved nest data for aerial surveys and ground line transects
                # and reconnaissance walks
                for (k in 1:K)
                        {
                        logit(rhonest[i,k,t]) <- zeta
                        zi[i,k,t] <- O.nest[i,t]*rhonest[i,k,t]
                        Z.nest[i,k,t] ~ dbern(zi[i,k,t])  # Occurrence data
                        }
                }
        }
}"
```

```
# INPUT DATA
# ------------------------------------------------------------------------------------------
# PARAMETERS:
#      G = total number of grid cell (1 km resolution)
#      T = total time period (3)
#      K = total number of sub-cell within grid cell (K=25 for subgrid size 200 m)
#      NEIGHBOUR[i,j] = neighbor indices for grid cell i
#      TOTNEIGH[i] = total number of neighbors for grid cell i
# ------------------------------------------------------------------------------------------
# DATA:
#      Y[i,j,t] = nest count data for grid cell i, time period t, and survey type
#             (aerial surveys and ground line transects)
#      Znest[i,k,t] = nest occurrence data from aerial and ground line transects and
#             reconnaissance walk surveys for sub-cell k, grid cell i, and time period t
# ------------------------------------------------------------------------------------------
# PREDICTORS:
#      ALT[i] = altitude at grid cell i
#      TEMP[i,t] = mean annual temperature at grid cell i and time period t
#      TEMP2[i,t] = quadratic value of mean annual temperature at grid cell i and time
#             period t
#      FOREST[i,t] = forest cover at grid cell i and time period t
#      MGV[i,t] = location on mangrove forest (binary) at grid cell i and time period t
#      PT[i,t] = location on peat forest (binary) at grid cell i and time period t
#      LOWL[i,t] = location on lowland forest (binary) at grid cell i and time period t
#      MONT[i,t] = location on mantane forest (binary) at grid cell i and time period t
#      FRGM[i,t] = location on fragmented forest (binary) at grid cell i and time period t
# ------------------------------------------------------------------------------------------
```
